# Maize diterpenoid sensing via Ste3 a-pheromone receptor and rapid germination of *Colletotrichum graminicola* oval conidia facilitating root infection

**DOI:** 10.1101/2024.04.05.588234

**Authors:** A. Y. Rudolph, C. Schunke, C. Sasse, L. Antelo, J. Gerke, G. H. Braus, S. Pöggeler, D.E. Nordzieke

## Abstract

Most plant pathogenic microorganisms have evolved to infect distinct host tissues. The maize anthracnose fungus *Colletotrichum graminicola* is known for its ability to invade above-ground tissues with asexual falcate conidia. In addition, *C. graminicola* produces a second asexual spore type, oval conidia. This study investigates the specific adaptations that make oval conidia suitable for maize root infection, demonstrating that only oval conidia exhibit root pathogen characteristics. These include the ability to germinate in soil and grow chemotropically toward root-secreted molecules. High-performance liquid chromatography/mass spectrometry (HPLC/MS) analyses combined with biological assays indicate that diterpenoids, such as dihydrotanshinone I, are likely responsible for the chemical attraction of *C. graminicola*. Genetic analysis identified the a-pheromone receptor CgSte3 as responsible for diterpenoid perception by the fungal pathogen. In conclusion, the understanding of maize anthracnose disease must be expanded to include an elaborate root infection cycle by oval conidia.

## Introduction

The health of crop plants and fruits is affected by a variety of organisms, ranging from viruses and microorganisms (bacteria, oomycetes, fungi) to complex eukaryotes (nematodes, parasitic plants) (Strange and Scott, 2005). Among these, fungi are prominent pathogens of major crops, reducing agricultural yield and quality by destroying plants preharvest and their fruits postharvest (Fisher *et al*., 2012, Silva *et al*., 2019). Timely, precise, and sustainable crop protection strategies are essential, particularly as the burden of fungal plant diseases is expected to shift due to climate change (Chaloner *et al*., 2021, Zhang, 2018).

Fungal plant pathogens specialize in entering host plants through distinct tissues (roots, shoots, stems, leaves) or at specific developmental stages (flower, fruit). Depending on the infected tissue, adapted signal-sensing machineries and infection strategies are required. Foliar pathogens sense the plant surface via its physical and chemical properties, inducing germination and pathogenicity programs (Kou and Naqvi, 2016). In contrast, fungal root pathogens initiate infection before direct contact with the host, sensing nearby plants through root exudates that trigger germination and chemotropic growth towards plant-secreted signals (Okubara and Paulitz, 2005, Steinkellner *et al*., 2005, Narula *et al*., 2009, Vives-Peris *et al*., 2020). Known chemical signals for this directed growth include sugars, amino acids, organic acids, fatty acids, and plant-derived enzymes, guiding pathogens like *Fusarium oxysporum*, *Fusarium graminearum*, and *Verticillium dahliae* to host roots, where they colonize root surfaces and invade the host (Deacon and Donaldson, 1993, Donaldson and Deacon, 1993, Ruttledge and Nelson, 1997, Tyler, 2002, Turrà *et al*., 2015, Sridhar *et al*., 2020, Vangalis *et al*., 2023), where they start root surface colonization and host invasion (Lagopodi *et al*., 2002, Zhao *et al*., 2016). Additionally, some fungi, such as *Magnaporthe oryzae*, primarily leaf pathogens, can infect root tissues under laboratory conditions by direct root treatment with conidia suspensions or mycelial plugs (Marcel *et al*., 2010, Dufresne and Osbourn, 2001).

The hemibiotrophic maize pathogen *Colletotrichum graminicola* is primarily known for infecting above-ground tissues like leaves and stems (Bergstrom and Nicholson, 1999). Current analyses estimate that anthracnose stalk rot (ASR) alone causes a 10-20% annual loss in maize harvest worldwide (2021, Belisário *et al*., 2022). *C. graminicola* produces two types of asexual spores: oval and falcate conidia. Unlike micro- and macroconidia in other pathogens, both spore types are metabolically active and generated through different developmental processes in distinct plant tissues (Zhang *et al*., 2014, Ohara *et al*., 2004, Panaccione *et al*., 1989). Falcate conidia develop in acervuli on infected leaves, characterized by dark, spike-like hyphae surrounded by masses of falcate spores. Oval conidia, however, pinch off from hyphae in the plant’s vascular system and are stored in parenchyma cells (Belisário et al., 2022, Sukno *et al*., 2008). These differences influence the morphological and physiological properties of both spore types. Falcate conidia are single-nucleate and dormant due to the production of mycosporines, while oval conidia contain variable nuclei and lack secondary metabolites promoting dormancy, leading to distinct infection strategies (Panaccione et al., 1989, Nordzieke *et al*., 2019b, Nordzieke, 2022). These findings suggest unexplored but specific roles of both spore types in the maize anthracnose cycle.

This study provides the first comparative analysis of the root infection potentials of both *C. graminicola* spore types. Our analyses reveal that only oval conidia exhibit the characteristics of a fungal root pathogen, including the ability to germinate and propagate in soil and grow chemotropically towards maize root exudates, thus causing disease symptoms under natural conditions. Chemical analysis and biological assays strongly indicate that maize root-secreted diterpenoids act as attractants for chemotropic growth. The *C. graminicola* a-pheromone receptor Ste3 was identified as the diterpenoid sensor in this host detection system. Our results suggest that oval conidia are better adapted to root infection under natural field conditions, with soil germination being the critical differentiator between the spore types. This study is the first to describe how the generation of a second spore type enables a fungal plant pathogen to efficiently infect a host through multiple tissues.

## Results

### Root dipping in *C. graminicola* oval conidia induce a symptomless maize root infection and disease spreading into maize stems

*C. graminicola* is the causal agent of maize anthracnose and generates two types of asexual spores. Falcate conidia are more efficiently causing anthracnose leaf blight (ALB) (Nordzieke et al., 2019b) and had been described earlier to infect roots (Sukno et al., 2008). In this study, the general abilities of both spore types to colonize the root surface and to spread within the host was compared. A *C. graminicola* wildtype strain expressing a tdTomato-labelled histone marker, CgM2::RH2B (Nordzieke et al., 2019b), enabling the easy detection of the growing fungus on the surface of the root, was analyzed. CgM2::RH2B oval or falcate conidia suspensions were used as inoculum for the infection of 5 d old maize roots prior to incubation under humid conditions. The microscopic evaluation shows production of several fungal structures as previously observed for root infection with falcate conidia (Sukno et al., 2008). At 4 d post infection, production of acervuli, the asexual fruiting bodies, was observed for both conidia types (Figure 1). Additionally, development of typical root colonization structures like long, fast growing runner hyphae, hyphopodia for root penetration and dark-pigmented resting structures (microsclerotia) were observed upon oval and falcate conidia colonialization (Supplementary Figure 1).

**Figure 1:**
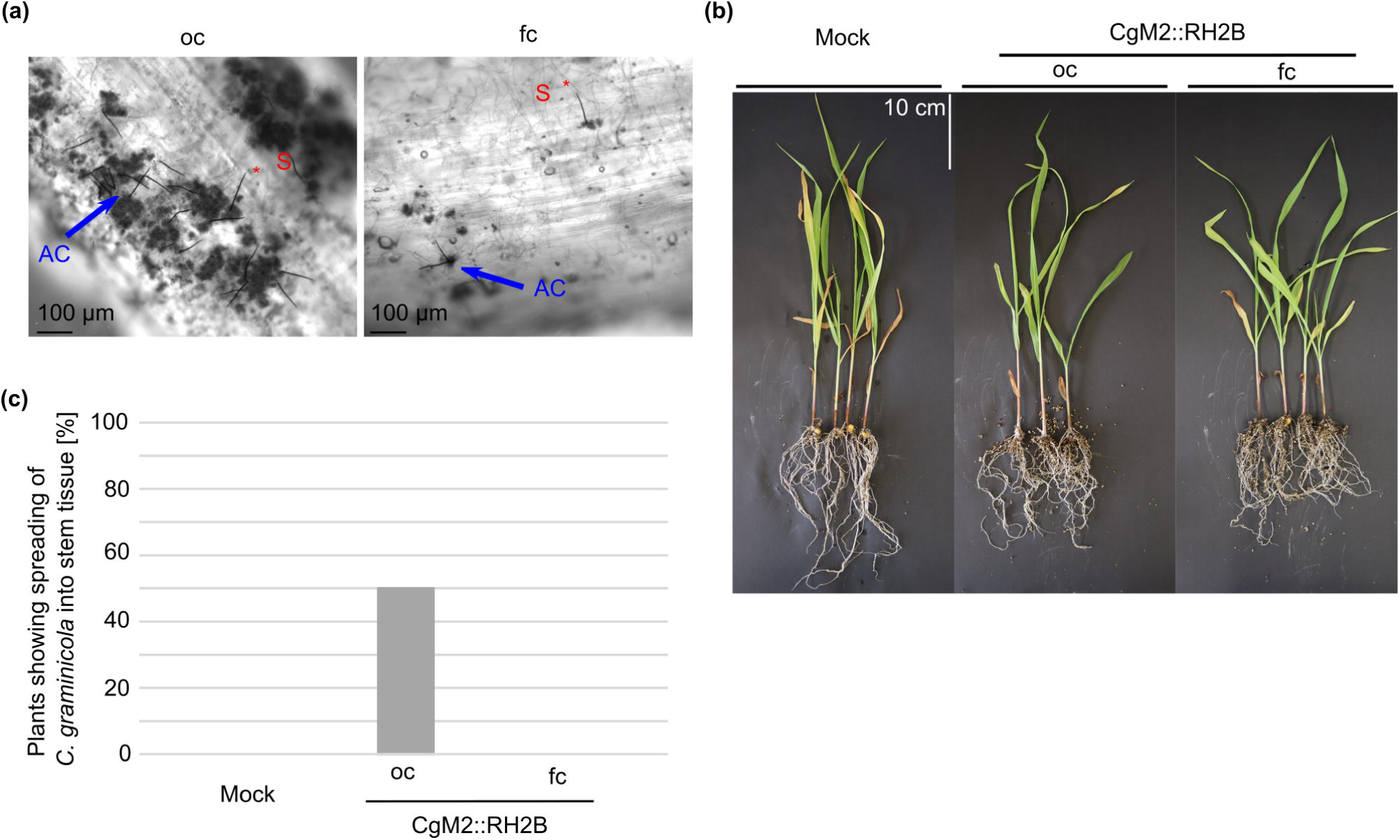
Symptom development after dipping root infection of *Z. mays* with *C. graminicola*. (a-c) Roots of 5 d old plants were dipped in oval (oc) or falcate (fc) conidial suspensions of CgM2::RH2B allowing for the detection of red-fluorescent nuclei during microscopic visualization. (a) *C. graminicola* development on roots 4 d after root dipping, AC = acervulus (blue arrow), S = setae (red asterisk), scale bar = 100 μm. (b-c) Symptom development 21 d after root dipping in conidial suspensions. Growth in vermiculite. (b) Representative depiction of plant development, scale bar = 10 cm. (c) Quantification of plants with spreading of *C. graminicola* in stem tissue by real-time PCR.

5 d old plants dipped in conidial suspensions were planted out in vermiculite to analyze symptom development on the growing plant. After 21 d of growth under humid conditions, length and biomass of above ground tissue was measured. As depicted in Supplementary Figure S2, no significant differences to mock inoculation were observed. A probable symptomless systemic infection was verified by taking stem samples as basis to detect fungal biomass in the respective plants in a real-time PCR approach. Specific detection of *C. graminicola* was ensured by the design of oligonucleotides binding in the species-specific region of ITS1 and ITS2 (Supplementary Figure 3). In 50% of plants infected with oval conidia fungal biomass was detected in stem material, whereas *C. graminicola* gDNA was absent in plants treated with falcate conidia (Figure 1 c). This supports a more effective *in planta* growth outgoing of oval conidia infection.

### Maize root exudate and diterpenoids elicit a strong chemotropic response by germlings derived from oval conidia

Directed growth to plant-secreted signals was described for several root-infecting pathogenic fungi (Nordzieke *et al*., 2019a, Turrà et al., 2015, Sridhar et al., 2020). The amount of germlings derived from oval and falcate conidia, which are able to redirect growth towards maize root-secreted signals, was quantified. A 3D printed device was applied for quantification, which was recently developed in our laboratory (Schunke *et al*., 2020). Oval conidia are well germinated after 6 h of incubation. They show typical localization of the polarity marker CgArp1-TagRFP-T (Groth *et al*., 2021) to the subapical region of growing hyphae. The corresponding hyphal tips point mainly to the gradient of maize root exudate (MRE), indicating active hyphal growth of the germlings to the root-secreted signals (Figure 2 a, b). In contrast, we were unable to detect any germinated falcate conidia, indicating that those spores are unable to germinate and redirect growth under the experimental conditions (Figure 2 a).

**Figure 2:**
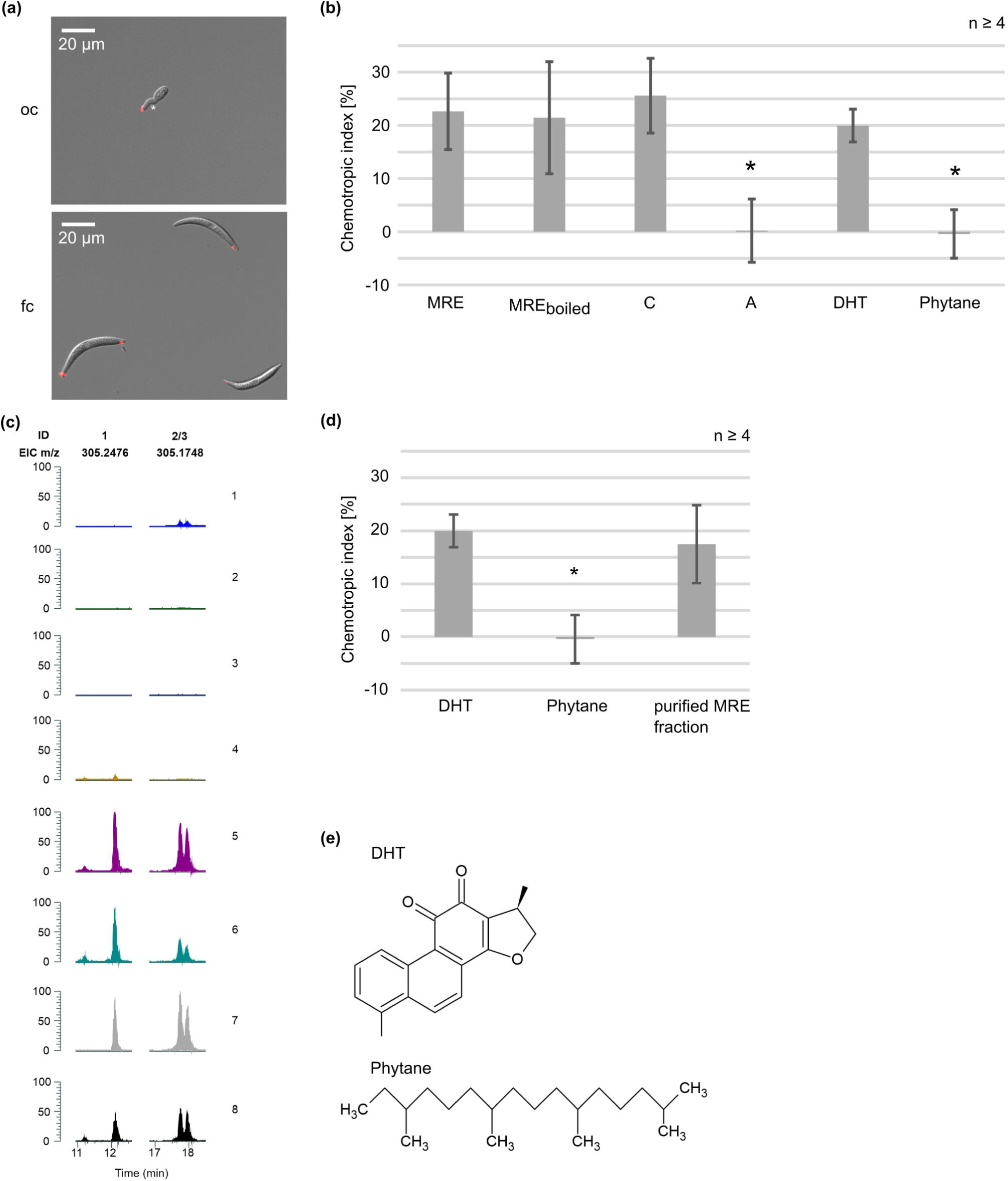
Chemotropic growth of *C. graminicola* conidia. (a) oval and falcate conidia of *C. graminicola* expressing red-labeled Arp1 (CgM2::Arp1-TagRFP-T, (Groth et al., 2021)) after 6 h of incubation facing a gradient of maize root exudate (MRE). Germination is indicated by a white asterisk, scale bar = 20 μm. (b) Chemotropic index of CgM2 oval conidia facing gradients of different chemoattractants including MRE and boiled MRE. Chloroform (C) and aqueous (A) phases gained from lipid extraction (chloroform:methanol) of MRE. Error bars represent SD calculated from n ≥ 4 experiments, * p < 0.05. (c) HPLC/MS analyses of non-attracting (fractions 1-4) and attracting (fractions 5-8) MRE fractions. Extracted Ion Chromatograms (EICs) for the identified metabolites 1 und 2/3. m/z 1 was detected in positive mode. ID (number of the compound in this study). The samples were extracted with ethyl acetate (1:1) or chloroform:methanol (1.8:2.2), n ≥ 2. Fractions: 1 = demineralized water; 2 = demineralized water used as basis for a chloroform:methanol extraction, chloroform phase; 3 = demineralized water used as basis for a chloroform:methanol extraction, aqueous phase; 4= MRE used as basis for chlorform:methanol extraction, aqueous phase; 5 = MRE after ethylacetate extraction; 6 = MRE boiled after ethyl acetate extraction; 7 = MRE used as basis for chloroform:methanol extraction, chloroform phase; 8 = MRE obtained from *C. graminicola* root-infected plants. (d) Chemotropic growth of *C. graminicola* oval conidia towards different chemoattractants. Error bars represent SD calculated from n ≥ 4 experiments, * p < 0.05, calculated with two-tailed *t*-test (e) Chemical structure of the tricyclic diterpenoid dihydrotanshinone I (DHT) and phytane, a linear diterpenoid. Chemical structures drawn with ACD/ChemSketch.

Next, we sought to identify the attracting molecule incorporated in MRE. Secreted peroxidases from the host were identified as chemoattractant signals in plant interaction studies with other phytopathogenic species (Turrà et al., 2015, Sridhar et al., 2020, Vangalis et al., 2023). Since the enzymatic activity of those enzymes are required for fungal attraction, boiling of tomato root exudate abolishes its attracting potential (Turrà et al., 2015, Nordzieke et al., 2019a). In contrast, boiling of MRE does not diminish the chemotropic growth index of *C. graminicola* germlings (Figure 2 b), indicating that peroxidases are not the attractant secreted from maize roots. Chloroform / methanol extractions were performed to explore further chemical properties of the active MRE, resulting in two phases with different chemical properties. Intriguingly, the chloroform phase, incorporating unipolar compounds such as lipids or lipid-associated molecules, induced a strong chemotropic response after solvent evaporation and resuspension in water. No directed growth was induced by the water phase typically containing nucleic acids (Figure 2 b). Attracting (MRE, boiled MRE, MRE chloroform phase of lipid extraction, MRE obtained from *C. graminicola* infected plants) and not attracting (demineralized water and its chloroform and aqueous phase of lipid extraction, MRE aqueous phase of lipid extraction) samples were compared using HPLC / MS analyses. Exudates generated from previously root-infected plants were also included, to test whether previous biotic stress by *C. graminicola* root infection might change the chemical composition of the MRE. Three molecules in the active fractions were consistently detected in three independent experiments, at the same time absent from the negative controls (Figure 2 c, Supplementary Figure 4). Interestingly, the double peak eluting at approximately 18 min was absent in the third replicate, except for the MRE sample from *C. graminicola* root infected plants. This indicates a flexibility of MRE composition dependent on the defense status of the plant. Comparing UV and MS2 spectra of the chemical compound released at 12 min (mass: 305,2476) with known diterpenoids from maize, we found the colorless 3β,15,16-trihydroxydolabrene (trihydroxydolabrene, THD) having a very similar MS2 spectrum (Supplementary Figure 5) (Mafu *et al*., 2018). After fractionation of this single peak, we verified its attracting potential in a chemotropic assay (Figure 2 d). We conclude it as very likely that the maize root-derived diterpenoid THD is attracting *C. graminicola* oval conidia to its host. In order to test non-maize diterpenoids for their attracting potential, we chose dihydrotanshinone I (DHT, originated from *Salvia* species), a cyclic diterpenoid like THD, and phytane, a linear diterpenoid. As depicted in Figure 2 d-e, oval conidia of *C. graminicola* are attraced by DHT. In contrast, the linear diterpenoid phytane did not induce a re-direction of *C. graminicola* germlings.

### The CgSte3 a-pheromone receptor is responsible for diterpenoid sensing in *C. graminicola*

In different *Fusarium* species, the pheromone receptors Ste2 (α-pheromone receptor) and Ste3 (a-pheromone receptor) activate the downstream Cell Wall Integrity (CWI) MAPK cascade, resulting in the re-direction of growth towards plant-derived peroxidases (Turrà et al., 2015, Sridhar et al., 2020, Sharma *et al*., 2022, Ramaswe *et al*., 2024). In contrast to other filamentous fungi, *C. graminicola* does not encode a *ste2* gene and the corresponding α-pheromone is likewise missing in the genome (Wilson *et al*., 2021). Therefore, a *Cgste3* deletion mutant was generated and characterized to test whether homologous pathway components are responsible for diterpenoid sensing by *C. graminicola* (Supplementary Figures 6 and 7). Intriguingly, germlings derived from ΔCgste3 oval conidia are unable to re-direct their growth to MRE and DHT (Figure 3 a), providing evidence that this receptor is required for the sensing of diterpenoid plant signals by *C. graminicola*.

**Figure 3:**
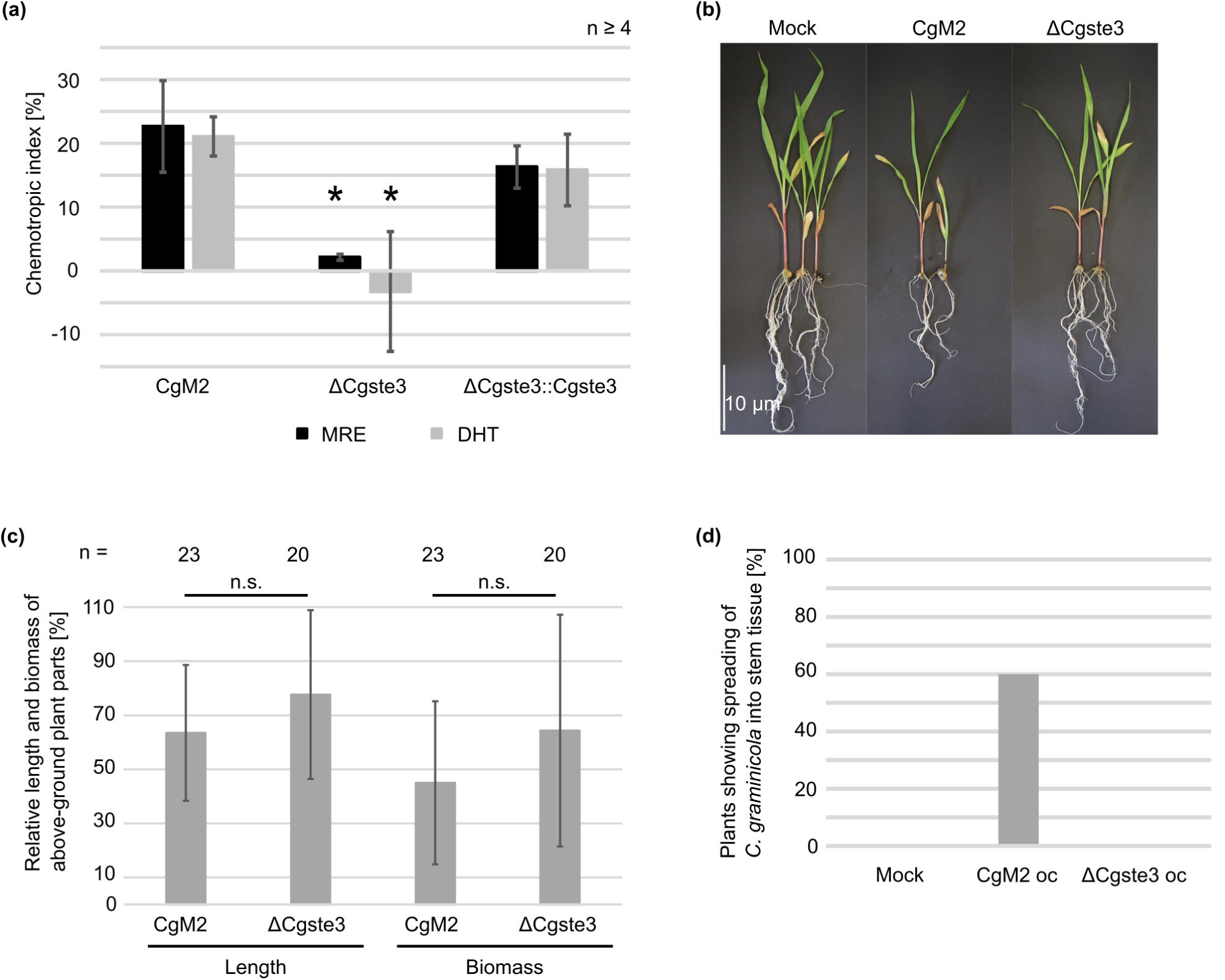
Perception of maize diterpenoids by a conserved fungal signaling pathway. (a) Chemotropic index of CgM2, Cgste3, and Cgste3::Cgste3 oval conidia facing gradients of different chemoattractants including MRE and dihydrotanshinone I (DHT). Statistical significance was calculated with CgM2 as reference. (b-c) Symptom development of plants co-incubated with conidia in vermiculite. (b) Representative depiction of plant development, scale bar = 10 cm. (c) Quantification of length and biomass of above-ground parts. Error bars represent SD calculated from n ≥ 20 experiments, ns = not significant. Statistical significance was calculated with two-tailed *t*-tests (d) Quantification of plants with spreading of *C. graminicola* in stem tissue with real-time PCR.

As many studies reported, spores of plant pathogenic fungi are adapted to germinate in the presence of an adequate host and to direct their growth towards its roots (Steinkellner et al., 2005, Li *et al*., 2013, Eze, 2010, Vangalis et al., 2023, Turrà et al., 2015). Maize seeds were disseminated in oval conidia-enriched soil of the *C. graminicola* wildtype strain CgM2 to mimic this natural infection situation under laboratory conditions (Figure 3). After three weeks of co-incubation, length and biomass of green plant parts were examined (Figure 3 b-c). In contrast to the results obtained by root-dipping, oval conidia co-incubated maize plants are significantly stunted and reduced in biomass compared to mock treatment and on some leaves typical anthracnose symptoms occurred (Supplementary Figure 8). Experiments outgoing from spore-enriched soil were also conducted with the mutant strain of ΔCgste3 (Figure 3 c-d). However, no difference between ΔCgste3 infections to wildtype was observed. To compare *in planta* growth between the wildtype and mutant strain, stem samples were used to detect of fungal biomass in a real-time PCR approach as described above. Whereas fungal DNA was present in 50% of plants co-incubated with oval conidia of the wildtype, oval conidia of the *Cgste3* deletion mutant did not spread into stem tissue (Figure 3 d).

### Successful germination of *C. graminicola* conidia in soil determines the ability to infect maize roots

Besides the ability to re-direct growth to MRE, one of the most prominent differences between oval and falcate conidia are the specific germination patterns. Germination of falcate conidia is regulated by a quorum sensing mechanism based on the secretion of mycosporines. In contrast, oval conidia do not secrete mycosporine-derivatives and thus germinate rapidly even when nutrients are absent (Nordzieke et al., 2019b). These spore-type specific adaptations suggested that there might be divergent germination patterns in vermiculite or other plant substrates, which influence the abilities of both spore types to infect the host plant’s roots.

The germination ability of oval and falcate conidia in soil of different compositions was analyzed to address this hypothesis (Figure 4 a, b). After incubation, the spore-contaminated soil was washed with a defined amount of water. The number of not germinated spores washed out was determined, assuming that germinating conidia would stick to the pieces of substrate. Independent of the substrate used, the fraction of non-germinated oval conidia dropped down already after 6 h of incubation (vermiculite: 75% reduction; vermiculite soil mix: 43.8% reduction). The number of oval conidia increased within the first two days of inoculation in both substrates. Although this effect was more pronounced in the vermiculite soil mix, a probable formation of new spores in the watered substrates is indicated. The main fractions of conidia were germinated in both spore samples after 10 d of incubation (vermiculite: 62% reduction; vermiculite soil mix: 92.8% reduction). In contrast, for falcate conidia the amount of not germinated spores detected remained stable, also after prolonged incubation for 10 d.

**Figure 4:**
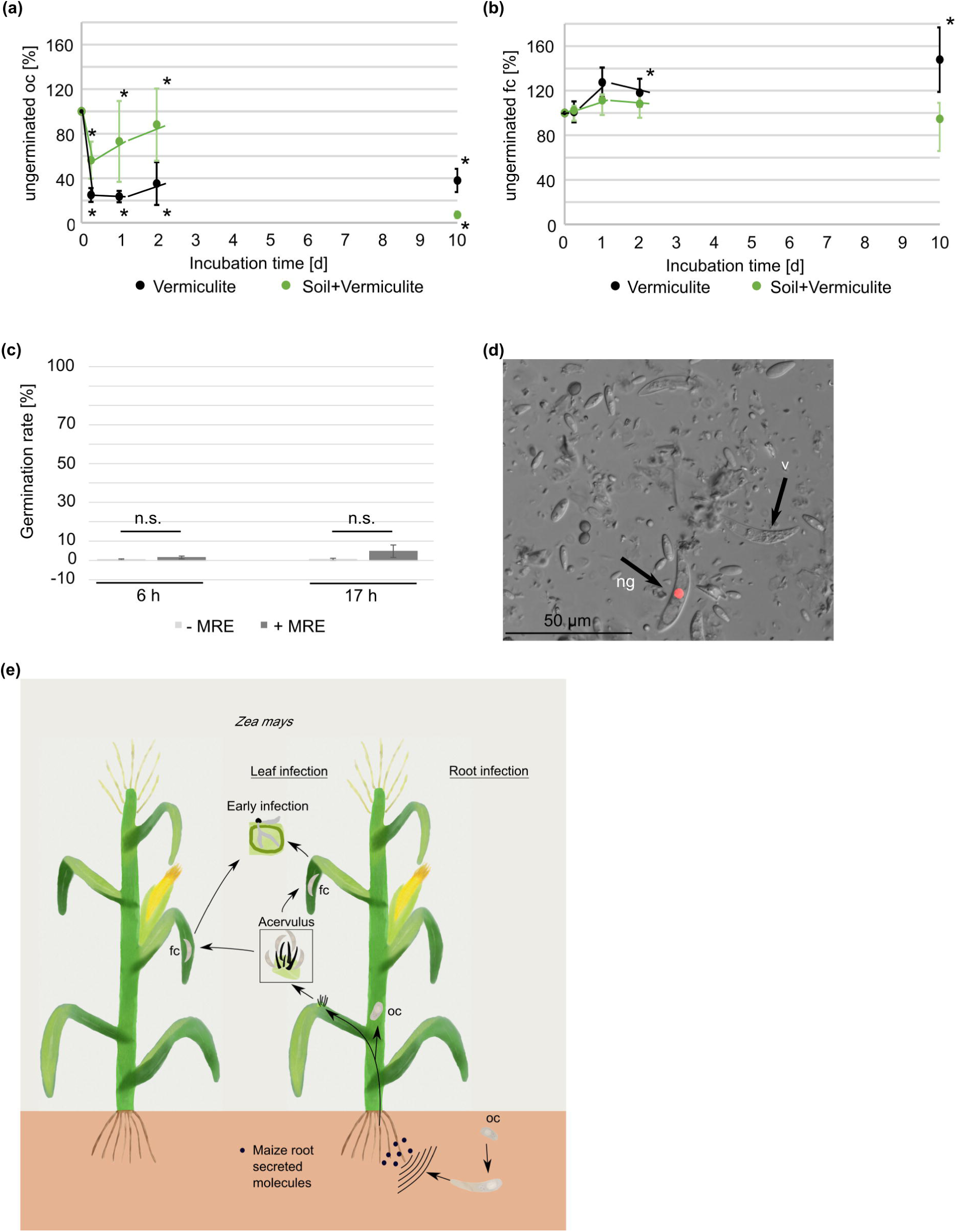
Germination of *C. graminicola* conidia. (a, b) Germination of oval (oc) (a) and falcate (fc) (b) conidia in vermiculite and a mixture of soil and vermiculite (4+1). Quantification of ungerminated conidia after incubation of 0 h, 6 h, 24 h, 48 h, 10 d in a Neubauer counting chamber. (c) Germination of falcate conidia (fc) on water agar after 6 h and 17 h with and without the supplement of MRE. Two-tailed t-tests were performed. * p < 0.05, n.s = not significant. (d) Falcate conidia (black arrows) in soil after a 10 d co-incubation with maize roots in vermiculite, ng = not germinated, scale bar = 50 μm (e) Model of the maize infection cycle. In the monocyclic part, oc of *C. graminicola* germinate in soil and grow chemotropically towards plant-secreted molecules. The fungus penetrates roots and systemically infects the plant leading to fc production in acervuli on leaves. Spreading of fc to nearby plants starts the polycyclic disease cycle. Leaf infection starts with inoculum and adhesion of spores, followed by development of the penetration structures.

It was shown for several root-interacting fungi that root exudate could induce germination (Steinkellner et al., 2005, Li et al., 2013, Eze, 2010). We thus tested whether MRE is able to induce germination of falcate conidia on axenic culture or in soil. For the first experiments, defined drops of falcate conidia were incubated on water agar with or without the addition of MRE. As depicted in Figure 4 c, MRE was unable to induce germination in falcate conidia. We further analyzed the effect of living maize plants on the germination ability. Consistently to the previous results, only non-germinated falcate conidia were detectable after 10 d of plant-co-incubation (Figure 4 d). Furthermore, many highly vacuolated and dead falcate conidia were detected, indicating that this spore type is unable to survive for a longer time in soil (Supplementary Figure S9).

## Discussion

The phytopathogenic fungus *Colletotrichum graminicola* produces two distinct types of asexual spores. The main finding of this study is that only oval conidia are adapted for infecting maize roots, whereas the falcate asexual spores invade above-ground tissues. This differentiation arises from two major factors: only oval conidia have the ability to germinate rapidly in soil and can efficiently sense maize root exudates, providing growth orientation towards the host. These findings provide new insights into the complex maize anthracnose cycle and have important implications for future crop protection strategies (Figure 4e).

Exudates from plant roots play several critical roles in plant health, such as adapting soil pH to enhance nutrient uptake, chelating toxic compounds, attracting beneficial microorganisms, and releasing antimicrobial substances to defend against pathogens (Vives-Peris et al., 2020, Zhang *et al*., 2020). During pathogen attacks, plants further adapt the composition of root exudates by synthesizing new antimicrobial compounds (Lanoue *et al*., 2010). Prominent examples of such antimicrobial compounds include class III peroxidases, which are secreted from plant roots and participate in reactive oxygen species defense pathways (Mika *et al*., 2010, Bindschedler *et al*., 2006, Wen *et al*., 2007). However, several pathogenic fungi can sense these peroxidases and, rather than being harmed or repelled, use them to guide themselves towards nearby host plants through a sophisticated amplification and sensing system involving fungal NADPH oxidase and pheromone receptors (Nordzieke et al., 2019a, Turrà et al., 2015, Sridhar et al., 2020, Vangalis et al., 2023).

In this study, we describe for the first time the ability of *C. graminicola* oval conidia to sense maize root exudates, provoking a robust chemotropic response. Chemical analysis and fractionation experiments revealed that the active substances secreted from maize roots are heat-stable and soluble in chloroform (Figure 2). Comparisons of the masses obtained from HPLC/MS analysis with known secondary metabolites showed a high similarity with diterpenoids. UV and MS2 spectra of one compound matched well with 3β,15,16-trihydroxydolabrene (THD). This colorless diterpenoid, secreted from maize roots, has antimicrobial properties (Mafu et al., 2018). Diterpenoids are involved in terpenoid-based defenses against various maize pathogens, including fungi such as *Rhizopus microsporus*, *Ustilago maydis*, and *F. graminearum*, as well as herbivorous and insect species (Zhou *et al*., 2023). Similar to class III peroxidases, the biosynthesis of several diterpenoids is induced by plant pathogens and defense-related phytohormones such as jasmonic acid and ethylene (Schmelz *et al*., 2011, Hiraga *et al*., 2001, Kidwai *et al*., 2020, Huffaker *et al*., 2011).

Our results show that diterpenoid and maize root exudate (MRE) perception in *C. graminicola* is mediated by the a-pheromone receptor Ste3, indicating a certain level of conservation in the sensing of host-secreted molecules despite the different chemical properties of the attractants (Turrà et al., 2015, Sridhar et al., 2020, Vangalis et al., 2023). However, unlike *Fusarium* and *Verticillium* species, the absence of the α-pheromone receptor Ste2 (Wilson et al., 2021) does not impede the perception of maize root secreted signals, suggesting differences in the perception process despite their similarities. These results indicate that the ability to use plant defense molecules for host detection has evolved multiple times and can adapt to different chemical compounds depending on the fungal-plant pathosystem involved.

A critical question is the relevance of chemotropic sensing by *C. graminicola* oval conidia and derived germlings for adapting this spore type for maize root infection. Our experimental data suggest that chemotropic sensing of maize root exudates is not essential for the *C. graminicola* root infection process per se but may have a variable impact depending on the spore concentration near the young maize plant. In a high inoculum scenario, exposure to germinated oval conidia might suffice to initiate root infection, although the process might be delayed, leading to slower spreading of *C. graminicola* within the host. Under field conditions, however, low spore inocula of one species likely make chemotropic guidance crucial for the pathogen to locate its host.

This study emphasizes the importance of proper germination for plant infection Falcate but not oval conidia generate mycosporines, which are self-inhibitors of germination produced at high spore densities, low availability of monosaccharides like glucose, and in the absence of a living plant surface (Leite and Nicholson, 1992, Nordzieke et al., 2019b). Although sugars are present as poly- and monosaccharides in soil (Gunina and Kuzyakov, 2015) our results demonstrate that these concentrations are insufficient to inhibit mycosporine generation and break the dormancy of falcate conidia. Despite identifying mycosporines as germination regulators for *C. graminicola* in the 1990s (Leite and Nicholson, 1992), the biosynthesis and regulation of these compounds remain unclear, warranting further studies to unveil this critical aspect of the maize anthracnose fungus.

In conclusion, our results indicate that oval and falcate conidia are adapted for distinct phases of the overall maize anthracnose cycle. Oval conidia act as primary inocula, initiating root infections through rapid germination in soil and chemotropic sensing of host roots. Following primary root infection, the maize plant becomes systemically infected, and acervuli develop on leaves, giving rise to falcate conidia. These falcate conidia are dispersed by wind and rain to other maize plants, spreading the disease through continuous leaf infections. To our knowledge, such specialization of two different asexual spore types for host infection has not been reported for any other plant pathogenic fungus. However, it could represent an unexplored adaptation strategy in many species and be highly relevant for effective crop protection.

## Experimental procedures

### Strains, growth conditions, and collection of spores

The sequenced *C. graminicola* (Ces.) G.W. Wilson (teleomorph *Glomerella graminicola* D. J. Politis (Forgey *et al*., 1978)) wildtype strain M2 (also referred to as M1.001) was used (O’Connell *et al*., 2012). Falcate conidia production of *C. graminicola* was induced by growth on oat meal agar (OMA; 1L contains 50 g ground oat flakes (Alnatura, Bioland)) and the exposition to red (660-665 nm) and blue (450-455 nm) light in a ratio 3:1 at 23°C (Nordzieke et al., 2019b). After 14-21 d collection of falcate conidia with 0.02% Tween 20 solution. For the generation of oval conidia, liquid complex medium containing sucrose (CMS, 1% glucose, 0.1% yeast extract, 0.1% peptone, 0.5 M sucrose, 5.3 mM Ca(NO_3_)_2_) 0.073 mM KH_2_PO_4_, 1.04 mM MgSO_4_, 0.46 mM NaCl) was inoculated with 50 µl falcate spore stock solution or mycelial plugs. After germination induction at 23°C for 2-3 d (80 rpm), further incubation took place for 5-8 d without agitation in darkness. Oval conidia are collected by filtration through Miracloth (Merck-Millipore, Darmstadt, Germany) followed by centrifugation and a single washing step with demineralized water.

### Generation of plasmids and *C. graminicola* strains

A ΔCgste3 deletion mutant and the complementation strain ΔCgste3::Cgste3 were generated to analyze whether the a-pheromone receptor CgSte3 is involved in the sensing of maize root exudates (MRE). Assembly of the corresponding plasmids was conducted using the NEBuilder HiFi DNA Assembly Cloning Kit (New England Biolabs) according to the instruction manual. All plasmids were verified using DNA hydrolysis and sequencing using appropriate enzymes and primers. Oligonucleotides, strains and plasmids used are listed in supplementary tables S1-S3.

As basis for homologous recombination in *C. graminicola*, the plasmid pCgste3_KO was generated by the assembly of three PCR fragments into pJet1.2 (Thermo Scientific). The fragments 5’ *Cgste3* (Ste3_P_fw//Ste3_P_rv; 1,043 bp) and 3’ *Cgste3* (Ste3_T_fw//Ste3_T_rv; 1031 bp) were amplified from *C. graminicola* genomic DNA. The *hph* cassette, mediating resistance to hygromycine B, was generated using the plasmid pRS-hyg as template (hph-f//hph-r; 1,417 bp) (Bloemendal *et al*., 2012).

For ΔCgste3 complementation, the plasmid pCgste3_nat was constructed. The 5’ and 3’ regions of *Cgste3* were amplified together with the *Cgste3* gene in a PCR using genomic DNA of *C. graminicola* as template (Ste3_c_fw//Ste3_c_rv; 3,215 bp). As backbone for the assembly reaction served pJet_nat linearized with *Eco*RV (Nordzieke, 2022), resulting in a resistance to nourseothricin-dihydrogen sulphate in *C. graminicola* strains transformed with this plasmid.

Prior to transformations in *C. graminicola*, the plasmids pCgste3_KO and pCgste3_nat were linearized using the enzymes *Nae*I and *Sca*I, respectively. Oval conidia of CgM2 (transformation of pCgste3_KO) or ΔCgste3 (transformation of pCgste3_nat) served as basis for the generation of protoplasts as described previously (Groth et al., 2021). After transformation, regenerating protoplasts were selected on medium containing hygromycin B (500 µg/ ml, transformation of pCgste3_KO) or nourseothricin-dihydrogen sulphate (70 µg/ ml, transformation of pCgste3_nat). To obtain homokaryotic strains, single spore isolations were performed of antibiotic-resistant and PCR-verified primary transformants (Nordzieke, 2022).

Single spore isolates of ΔCgste3 were verified by Southern Blot analyses. Hydrolysis of genomic DNA (gDNA) was performed with the enzyme *Nae*I (Thermo Scientific). For the visualization of successful deletion of the *Cgste3* gene via homologous recombination, the 3’ region of *Cgste3* was amplified in a PCR reaction (Ste3_T_fw//Ste3_T_rv; 1,031 bp) and used as specific probe in the following hybridization reaction (expected sizes: CgM2 2,196 bp, ΔCgste3 7,896 bp, Supplementary Figure S6). Re-integration of *Cgste3* into the ΔCgste3 deletion strain was tested via PCR using the primer pair Ste3_fw//Ste3_rv (1,136 bp), verifying ΔCgste3::Cgste3 primary transformants and single spore isolates (Supplementary Figure S7).

### Root infection analyses

Two different methods were applied for all experiments, to study different aspects of *C. graminicola* root infection of the *Zea mays* cultivar ‘Mikado’ (KWS SAAT SE, Einbeck, Germany). Unless otherwise stated, incubation of plants was performed in a PK 520 WLED plant chamber (Poly Klima Climatic Growth System, Freising, Germany) using a day/night cycle of 12 h 26°C/12 h 18°C. The strains CgM2, CgM2::RH2B, expressing tdTomato labeled histone 2B (Nordzieke et al., 2019b), and ΔCgste3 were used for root infection experiments.

### Infection by a root-dipping technique

*Z. mays* roots were dipped in conidial suspensions when 5 d old as described previously by Sukno et al. 2008 to examine the general ability of *C. graminicola* to infect roots (Sukno et al., 2008). Early (4 dpi) or late (21 dpi) developed symptoms were examined microscopically. First, seeds of *Z. mays* were surface sterilized in 10% sodiumhypochlorite for 10 min and incubated for 5 d in rectangle petri dishes (82.9923.422, Sarstedt, Nümbrecht, Germany) on wet blotting paper (BF2 580x 600mm, Sartorius, Göttingen, Germany). Incubation took place in darkness using standard plant growing conditions. The developed roots of 5 d old seedlings were soaked in solutions of either oval or falcate conidia of CgM2::RH2B (c= 10^7^ x ml^-1^) for 1h at 100 rpm. As mock control, roots were soaked in sterile demineralized water. For follow up microscopic examinations, the *C. graminicola* coated plants were incubated on wet blotting paper for 4 d. 2 ml of demineralized water was added on the blotting paper after 2 d to prevent drought stress induction. For analysis of symptom development after 21 d, seedlings were planted in 40 g vermiculite (Vermiculite Palabora, grain size 2-3 mm, Isola Vermiculite GmbH, Sprockhövel, Germany) after dipping. The pots were covered with disposal plastic bags and sealed (Sarstedt, Nümbrecht, Germany). After 21 d of incubation, length and biomass of the above-ground plants were determined, pictures of the symptom development were taken and spreading into stem material assessed with real-time PCR (2.3.3).

### Root infections outward of *C. graminicola* enriched vermiculite

Maize seeds were planted in 40 g of vermiculite (Vermiculite Palabora, grain size 2-3 mm, Isola Vermiculite GmbH, Sprockhövel, Germany) enriched with oval conidia of CgM2 or ΔCgste3 in a concentration of 7.5 x 10^4^ x ml^-^1, to simulate a root infection outgoing from *C. graminicola* conidia present in agricultural soil. As mock control, seeds were sown in vermiculite mixed with water. To ensure high humidity, the pots were sealed in disposal plastic bags (Sarstedt, Nümbrecht, Germany). 21 dpi, length and biomass of the above-ground plant was determined and stem samples taken for follow up real-time PCR (2.3.3).

### Identification of *C. graminicola in planta* via real-time PCR

We developed a PCR-based identification method, to monitor the spreading of *C. graminicola* from infected roots into plant stems. First, oligonucleotides (ITS_P4_fw and ITS_P9_rv) were designed to bind to *C. graminicola*-specific sequences of ITS1 and ITS2 flanking 5.8S rRNA. The primer pair amplifies a 339 base pair fragment on a template of *C. graminicola* gDNA. Importantly, no amplicons were generated using gDNA from the fungi *S. macrospora, Verticillium dahliae, Verticillium longisporum, Aspergillus fumigatus* and *Aspergillus nidulans* as template in verification PCRs (Supplementary Figure 3 a). Therefore, the primer pair was used in the real-time PCR on *C. graminicola*-infected plant material. To obtain template DNA, stems from infected plants obtained by the infection protocols described, were surface sterilized for 2 min in 10% sodium hypochlorite solution. From the lower part, 0.5 cm of stem were discarded. The following 2 cm of the plant stems were frozen in liquid nitrogen and stored at −80°C (Supplementary Figure S3 b). The collected stem samples were grinded in liquid nitrogen to fine powder. 500 µl extraction buffer (Tris HCl pH 8.5 10 ml (from 1 M stock solution) EDTA 2.5 ml (from 0.5 M stock solution) SDS 3.125 ml (from 8% stock solution) NaCl 6.25 ml (from 2 M stock solution) demineralized water) was added to each sample. At this step, the plasmid peGFP-Cgatg8_gen (0.05 ng/µl) was added as an external standard for DNA losses during gDNA extraction. Further extraction was achieved with 8 M potassium acetate and chloroform. The DNA was precipitated with ice-cold isopropanol at −20°C for 2 h to overnight. The resulting pellet was resuspended in 50 µl demineralized water plus 2 µl RNase. Real-time PCR was performed with the MESA GREEN qPCR MasterMix Plus for SYBR Assay (Eurogentec, Seraing, Belgium). For detection, a CFX Connect Real Time PCR Detection System (Bio-Rad Laboratories, Hercules, CA, USA) was used. The external standard plasmid peGFP-Cgatg8_gen served as template for amplification with the oligonucleotide pair GFP-f and GFP-r. Follow up DNA sequencing and BLAST analyses verified the identity of the obtained fragment outgoing from gDNA of the stem samples as originating from *C. graminicola* (Supplementary Figure 10).

### Microscopy

Light (differential interference contrast (DIC)) and fluorescence microscopy was performed with the Axiolmager M1 microscope (Zeiss, Jena, Germany). The Photometrix coolSNAP HQ camera (Roper Scientific, Photometrics, Tucson, AZ, USA) was used to capture images and processing was performed with the ZEISS ZEN software (version 2.3, Zeiss). Visualization of expressed red fluorescent CgM2::RH2B was achieved by a 49005 Chroma filter set (exciter ET545/30x, emitter ET620/60 and beam splitter T570LP). At least three biological replicates were performed for each experiment.

### Generation and chemical analysis of maize root exudate

Roots of four 9 d old maize plants grown in vermiculite (Vermiculite Palabora, grain size 2-3 mm, Isola Vermiculite GmbH, Sprockhövel, Germany) under standard conditions to generate MRE. The roots were rinsed and placed in 40 ml demineralized water. After incubation for further 5 d in a plant chamber, MRE was sterile filtrated with a 0.45 µm filter (86.1197, Sarstedt, Nümbrecht, Germany).

An HPLC/MS analysis was applied to analyze MRE for probable bioactive substances. The MRE was extracted with ethyl acetate (1:1) or chloroform:methanol (1.8:2:2). Prior to HPLC/MS analysis, the samples were solved in 500 µl methanol. The received suspensions were centrifuged at 13000 rpm for 10 min at 4°C to remove unsolved particles. 400 µl of the supernatants were transferred to LC-MS glass vials. A Q-Exactive^TM^ Focus orbitrap mass spectrometer coupled to a Dionex Ultimate 3000 HPLC (Thermo Fisher Scientific) was used for analysis. HPLC was carried out using a Acclaim^TM^ 120, C_18_, 5 µm, 120 Å, 4.6 x 100 mm. 5 µl of each sample was injected for analysis. Running phase was set as a linear gradient from 5 - 95% (v/v) acetonitrile/0.1 formic acid in 20 min, plus 10 min with 95% (v/v) acetonitrile/0.1 formic acid) with a flow rate of 0.8 ml/min at 30°C in addition. The measurements were performed in positive and negative modes with a mass range of 70-1050 m/z. UV/Vis spectra were recorded with a Thermo Scientific™ Dionex™ UltiMate™ 3000 Diode Array Detector. MS2 data were recorded at a HCD energy of 35 eV. For data analysis the software FreeStyle^TM^ 1.4 (Thermo Fisher Scientific) and the database “Dictionary of Natural Products” were used.

For the HPLC fractionation of MRE, the extraction of 8.5 L of exudate was performed with an equal amount of ethyl acetate. The solvent was evaporated in a rotary evaporator and the resulting extract (387 mg) applied onto a preparative HPLC-system. HPLC purification was performed on a Shimadzu LC system equipped with a LC-20AP pump, an FCV-200AL valve, a CBM-40 controller and an FRC-10A fraction collector. Separations were done on a preparative reversed phase C18 column (Waters Sunfire C18, particle size 5 µm, 19 x 250mm) and a semipreparative reversed phase C18 column (Waters Sunfire C18, particle size 5 µm, 10 x 250mm) at room temperature. A linear gradient starting from 1% 0.1% TFA and 99% Acetonitrile to 100% Acetonitrile in 20 min, then maintaining 100% Acetonitrile for 3 min and an additional equilibration time of 7 min was used at a flow rate of 17.1 ml/min. Injection volume of sample solution was 1 ml. Volume of collected fractions were 3 ml. A Shimadzu SPD-M40A diode array detector was used from 200-800 nm for recording of spectra and detection of separated secondary metabolites at 210, 250, 300, 350 and 400 nm. Data and spectra were analyzed using the software LabSolution 5.110. LC-ESI-MS analysis was performed on a Shimadzu LC system equipped with two LC-40D pumps, a DGU-405 degassing unit, a SIL 40C autosampler, a SCL-40 controller and a CTO-40C column oven. Mass Analysis was performed with a single quadrupole LCMS-2020 system. Separations were done on an analytical reversed phase C18 column (Macherey Nagel, Nucleoshell RP18, particle size 2.7 µm, 2 x 100mm) at 40°C. A gradient starting from 95% 0,1% FA and 5% Acetonitrile with a holding time of 2 min to 100% Acetonitrile in 6 min, then maintaining 100% Acetonitrile for 2 min and an additional equilibration time of 4 min was used at a flow rate of 0.3 ml/min. Injection volume of sample solution was 1 µl. A Shimadzu SPD-M40A diode array detector was used from 200-800 nm for recording of spectra and detection of secondary metabolites at 210 nm, 250 nm, 300 nm, 350 nm, 400 nm, 450 nm and 500 nm. A single quadrupole LCMS System-2020 (Shimadzu) with ESI was used with following parameters: Scan mode, positive and negative, starting from 150 to 1000 m/z with acquisition time from 2 to 10 min. DL temperature 250 °C, nebulizing gas flow 1.5 L/min, drying gas flow 10 L/min and the temperature of the heat block was 200 °C. Data and spectra were analyzed using the software LabSolution 5.109.

### Chemotropic assay

Quantification of chemotropic growth of different *C. graminicola* germlings derived from oval and falcate conidia, towards MRE, different MRE fractions, oryzalexin D, dihydrotanshinone I, phytane, and attractants within MRE, was performed after 6h of incubation using a 3D printed device as described previously (Schunke et al., 2020, Groth et al., 2021). From raw data, the chemotropic index was calculated, showing the difference of chemotropic attraction compared to non-stimulating conditions (Turrà et al., 2015).

### Analysis of *C. graminicola* germination patterns

For quantification of germination in soil, 1.875 ml spores solution (c = 1.5*10^6) of either oval or falcate conidia was mixed in 5 ml vermiculite or soil (I)-vermiculite (mix of soil (I) (Einheitserde Classic, Patzer Erden GmbH, Sinntal-Altengronau, Germany) and vermiculite (Vermiculite Palabora, grain size 2-3 mm, Isola Vermiculite GmbH, Sprockhövel, Germany), ratio 4+1) mixture. After incubation (0 h, 6 h, 24 h, 48 h, 10 d), 2.5 ml water was added to wash out remaining spores. To access the rate of remaining not-germinated spores, those were quantified in a Neubauer counting chamber. 50 µl spore solution (c = 10^6 x ml^-1^) was dropped on water agar (1L contains 10 g agarose, 10 g serva agar, 2.1 g NaNO_3_). After evaporation of the liquid, 20µl MRE was added on top of the spores to analyze the effect of MRE on germination. Microscopy of germination behavior of falcate conidia in soil was performed after 24 h incubation in either vermiculite, soil (I)-vermiculite (4+1) mixture or soil (II) -compost soil- sand mixture (mix of soil (II) (Einheitserde Type P 25, Archut Erdenwerk, Vechta, Germany; compost soil, steamed for 24 h at 105°C, botanical garden, Göttingen, Germany; sand, grain size 0-2mm, August Oppermann Kiesgewinnungs- und Vertriebs-GmbH, Northeim, Germany; ratio 3+3+1).

### Statistics

The T-test for unequal variances was used for all experiments in this study(Ruxton, 2006).

## Acknowledgements

We thank Gabriele Beyer, Gertrud Stahlhut and Christiane Grünewald for excellent technical assistance. Genomic DNA of *Verticillium dahliae* and *Verticillium longisporum* was provided by Rebekka Harting. Genomic DNA of *Aspergillus fumigatus* and *Aspergillus nidulans* was provided by Anja Strohdiek. This work was funded by the Deutsche Forschungsgemeinschaft (Bonn-Bad Godesberg). Grants were provided to DEN (project NO 1230/3-1 (project number 447175909 and to DEN and GHB (IRTG 2172: PRoTECT program (GRK 2172, project number 273134146)).

## Data availability statement

The data that support the findings of this study are available from the corresponding author upon reasonable request.

## Supporting information legends

**Supplementary Figure 1:**
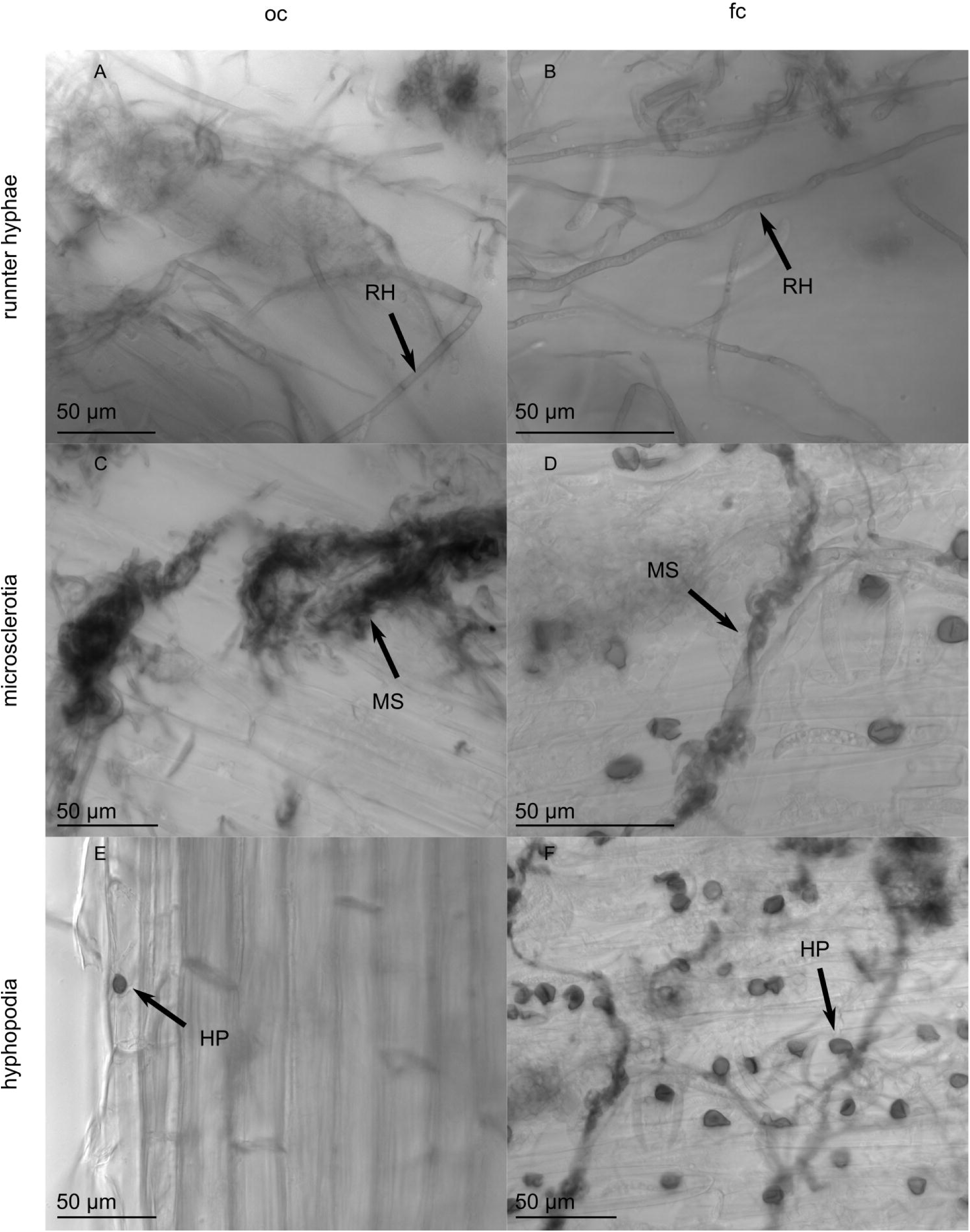
Microscopic assessment of *C. graminicola* colonizing and infecting roots of *Z. mays*. Roots of 5 d old maize plants were soaked in suspensions of oval (oc) or falcate (fc) conidia (c=10^7 x mL^-1^) and incubated on wet blotting paper for 4 d. RH= runner hyphae, MS = microsclerotia, HP = hyphopodia, scale bar = 50 μm.

**Supplementary Figure 2:**
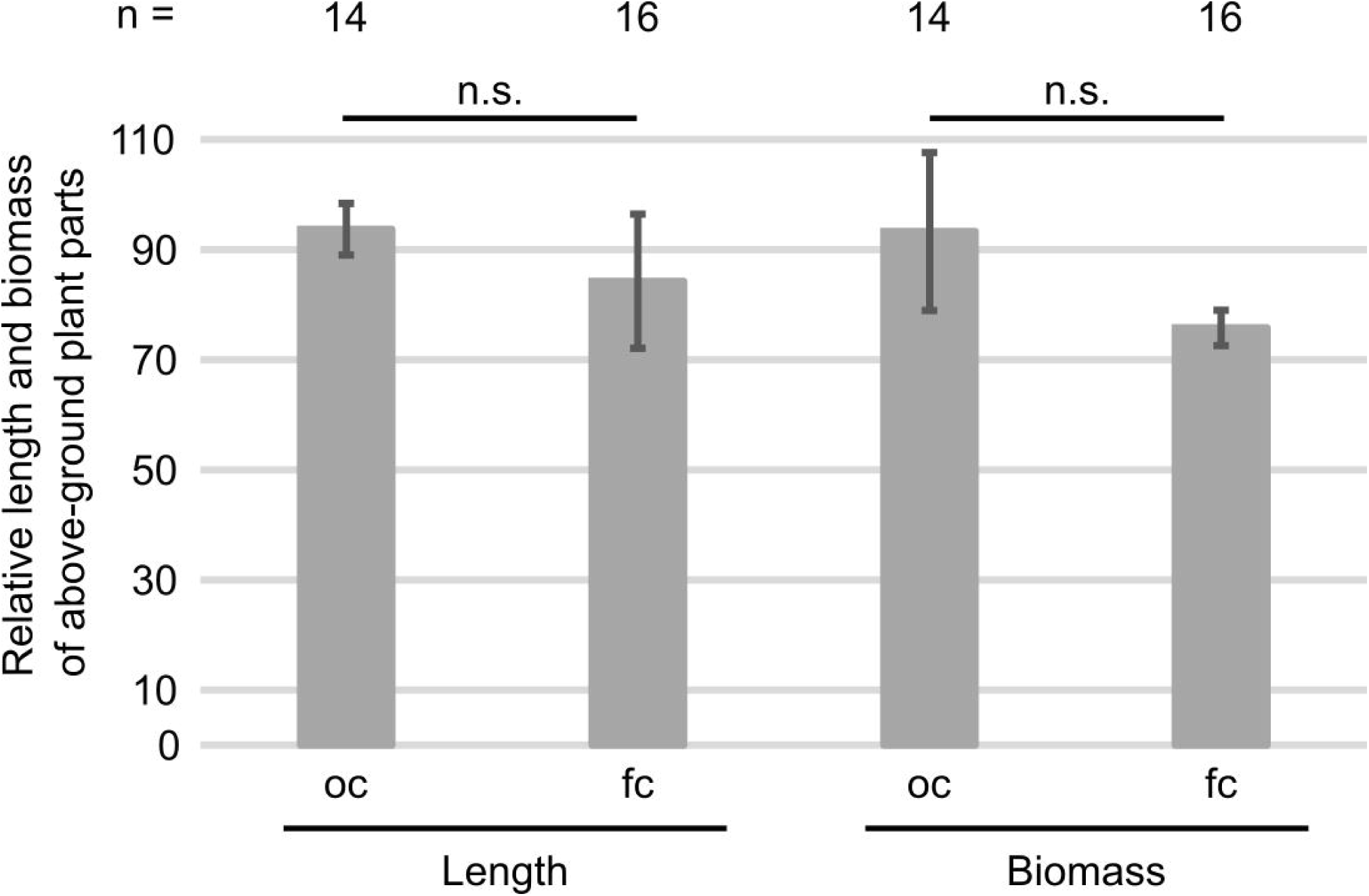
Symptom development after dipping root infection of *Z. mays* with *C. graminicola*. Roots of 5 d old plants were dipped in conidial suspensions. Subsequently, the plants were incubated in vermiculite for 21 d. Quantification of length and biomass of the above ground plant parts. Error bars represent SD calculated from n ≥ 14 experiments, ns = not significant.

**Supplementary Figure 3:**
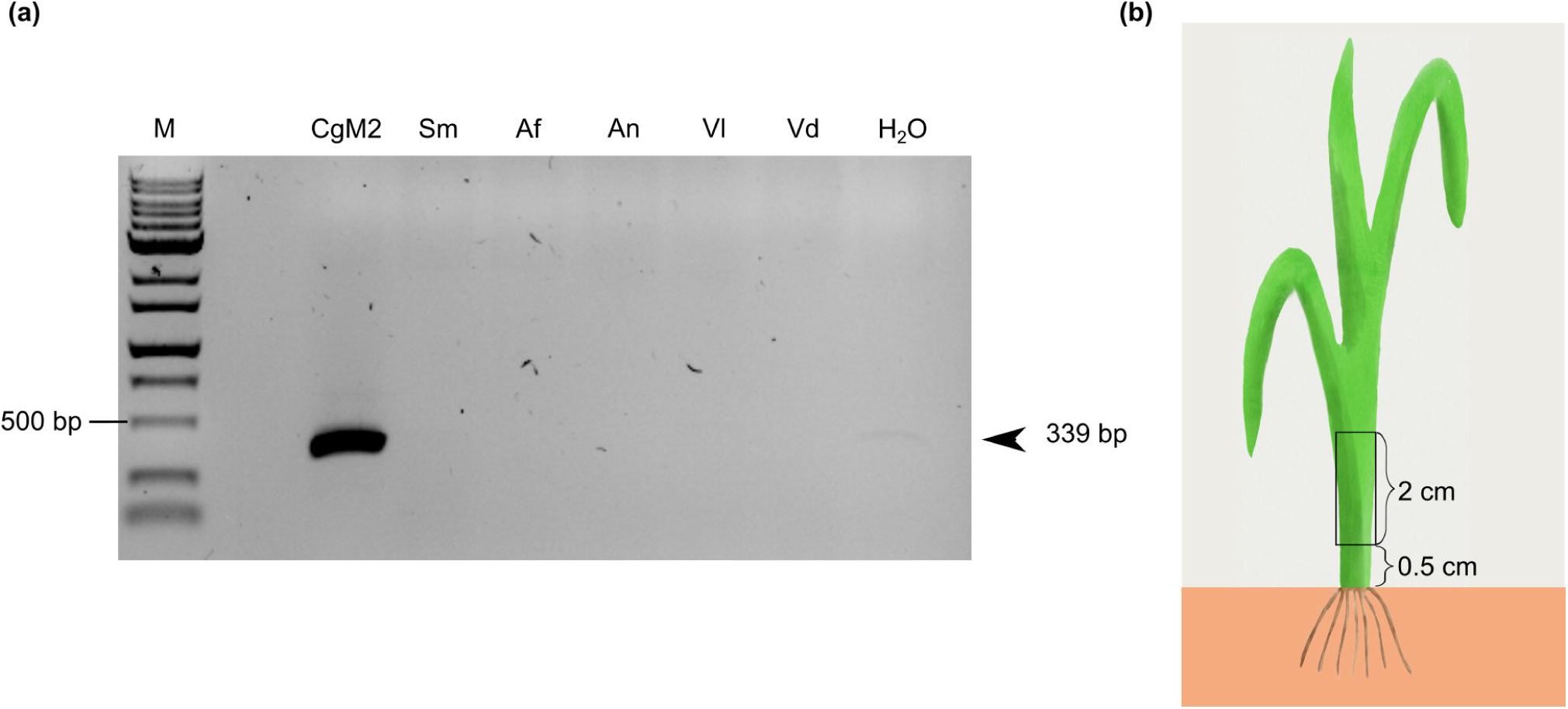
Establishment of the real-time PCR to quantify plants with spreading of *C. graminicola* in stem tissue. (a) Schematic depiction of sample collection. 0.5 cm above the kernel were discarded and gDNA extraction was performed from 2 cm stem above. (b) The primer ITS_P4_fw and ITS_P9_rv are designed to specifically bind CgM2 but no other fungi shown for *Sordaria macrospora* (Sm), *Aspergillus fumigatus* (Af), *Aspergillus nidulans* (An), *Verticillium longisporum* (Vl) and *Verticillium dahliae* (Vd). bp = base pairs.

**Supplementary Figure 4:**
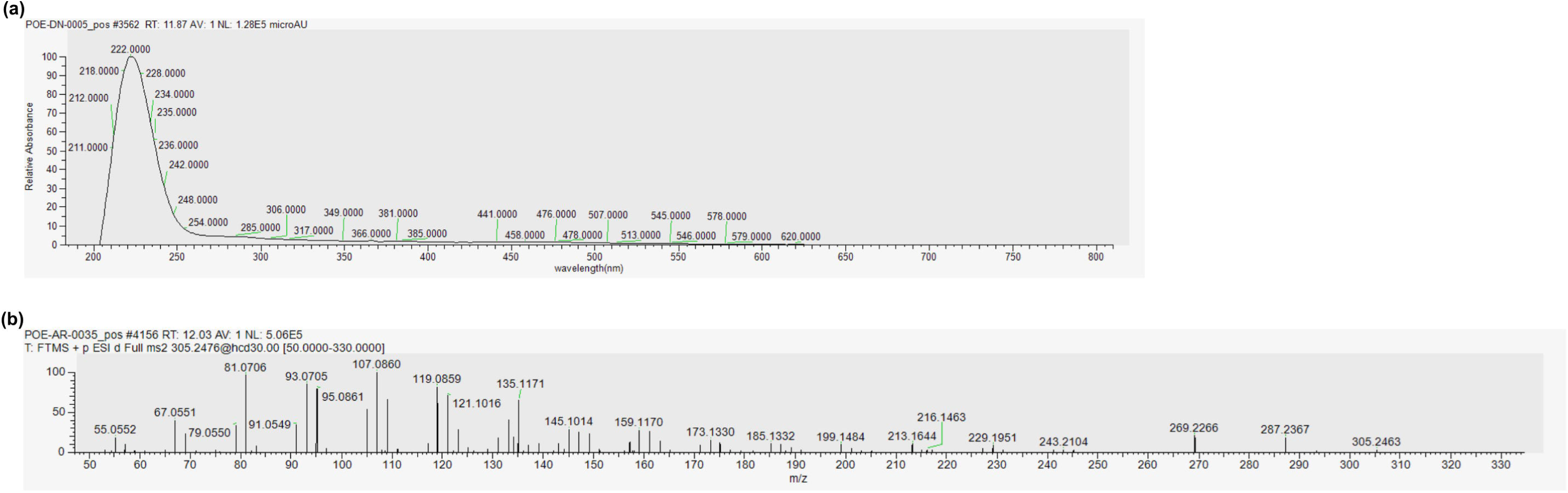
Chromatogram of HPLC/MS analysis of maize root exudate samples. (A-D) non-attracting samples (A) Demineralized water. (B-D) Lipid extraction samples. (B) Chloroform phase of demineralized water. (C) Aqueous phase of demineralized water. (D) Aqueous phase of MRE. (E-H) attracting samples. (E) MRE. (F) Boiled MRE. (G) Lipid extraction chloroform phase of MRE. (H) MRE of plants co-incubated with oval conidia of CgM2.

**Supplementary Figure 5:**
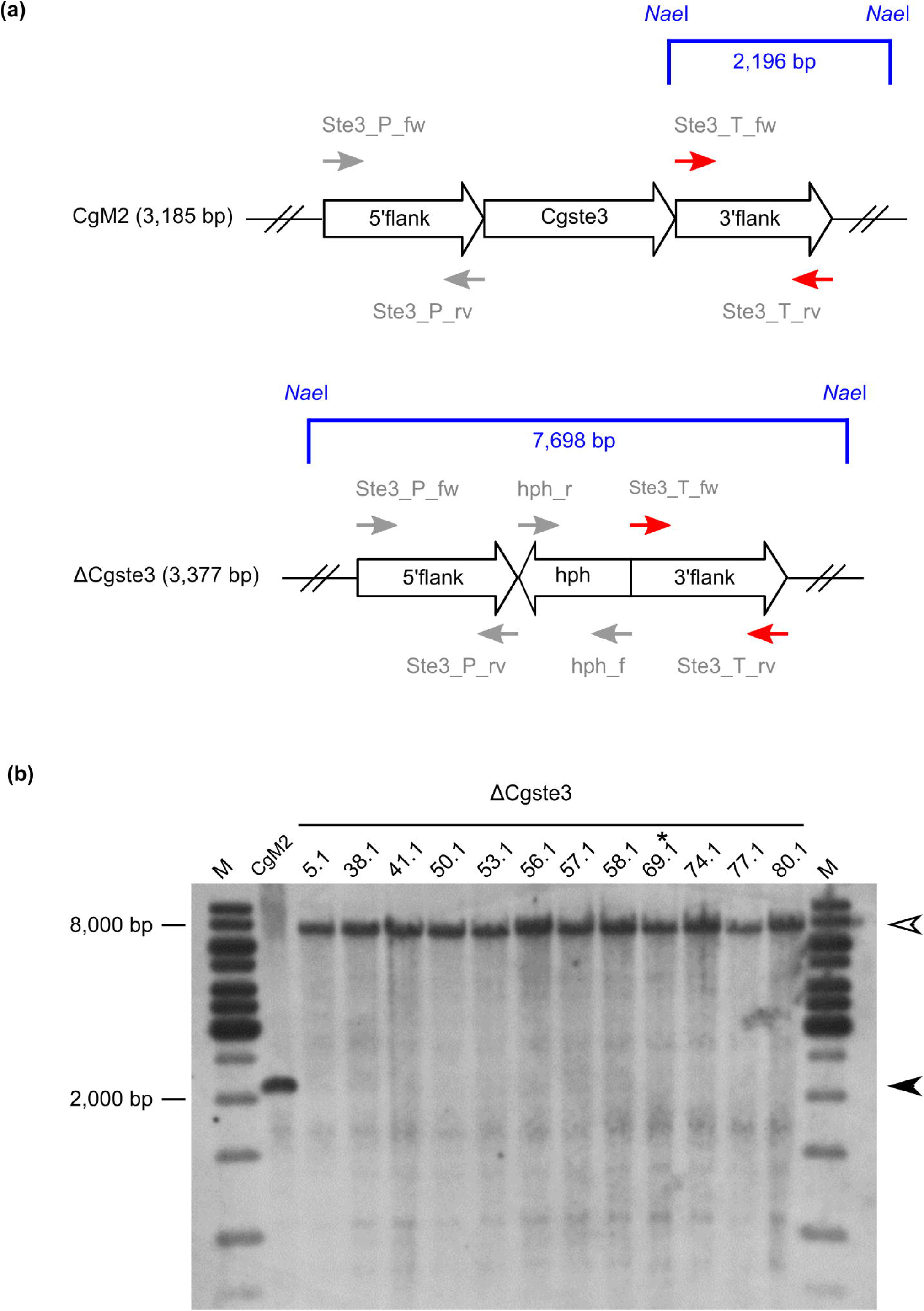
UV and MS2 spectra of compound 1, eluting at 12 min from HPLC. UV/Vis spectra were recorded with a Thermo ScientificTm DionexTm Ultim ateTm 3000 Diode Array Detector. MS2 data were recorded at a HCD energy of 35 eV.

**Supplementary Figure 6:**
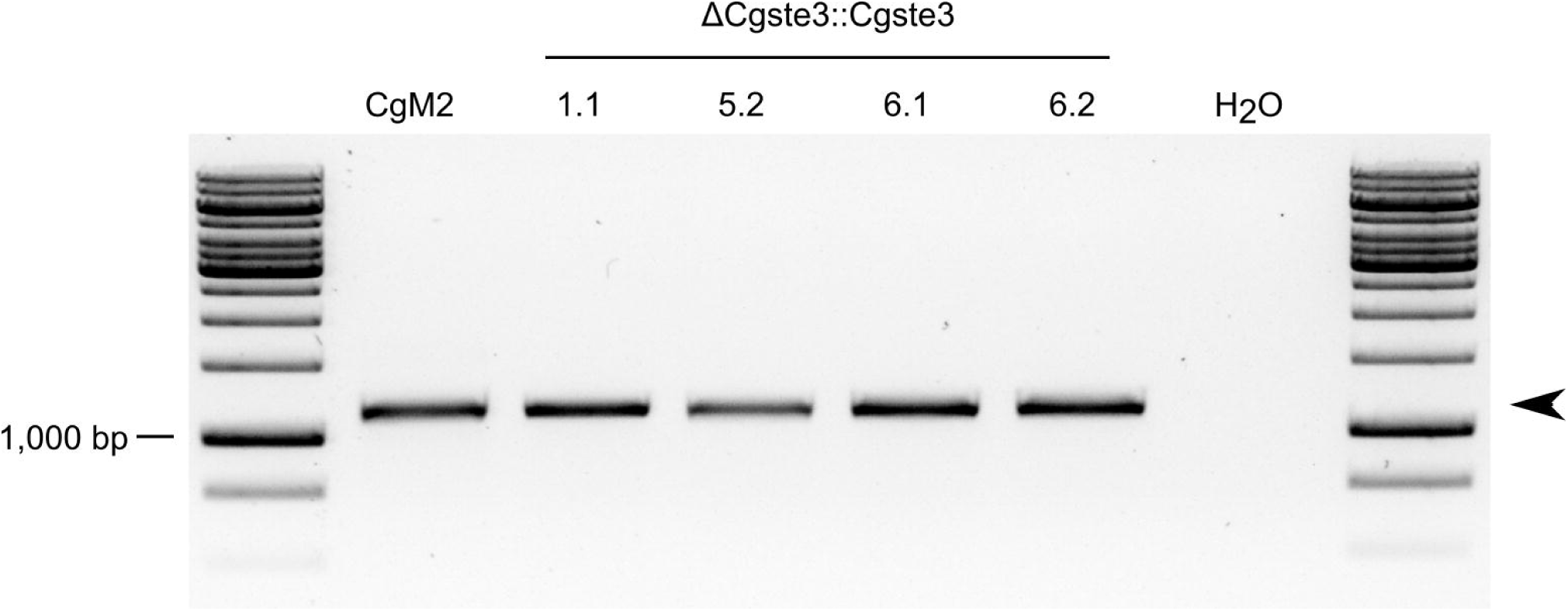
Generation of a *Cgste3* deletion strain in *C. graminicola*. (a) Strategy for ΔCgste3 generation. Genomic loci of *Cgste3* in CgM2 wildtype and deletion strain. Primer binding sites for the generation of the deletion construct are indicated, arrows indicate the amplification direction, red arrows indicate the oligonucleotides used for the generation of a *Cgste3*-specific probe for southern blot analysis. Recognition sites for *Nae*I and the expected band sizes are depicted in blue. (b) Verification of homologous integration of a *hph*-resistance cassette into the *Cgste3* locus via Southern Blot hybridization. Expected band sizes after hydrolysis of gDNA with *Nae*I are indicated with black (CgM2: 2,196 bp) and white arrowheads (ΔCgste3: 7,896 bp). Strains used for phenotypic characterization are written in bold letters, the strain used for complementation is additionally indicated with an asterisk.

**Supplementary Figure 7:**
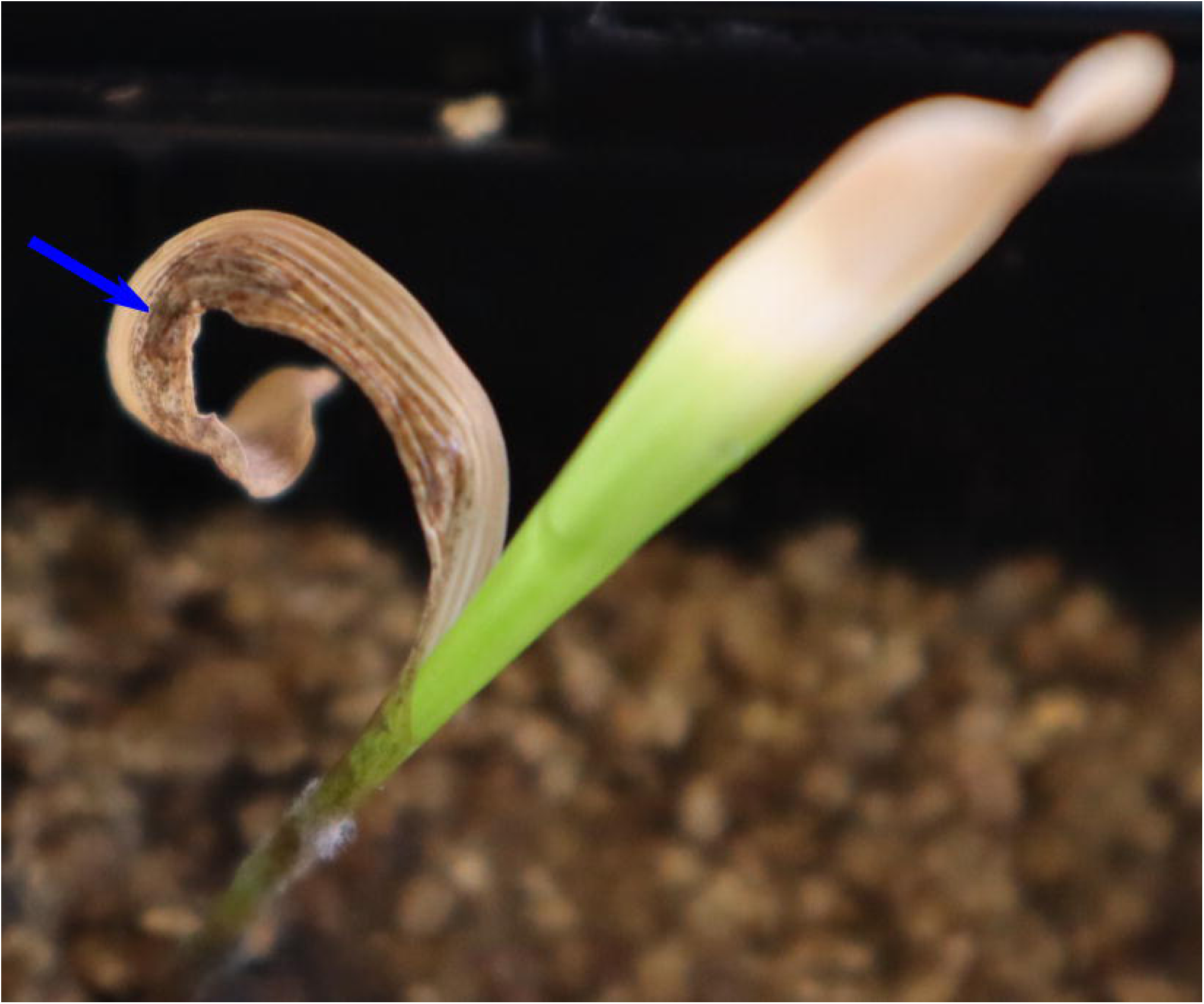
Verification of ΔCgste3::Cgste3. Amplification with the oligonucleotides Ste3_fw and Ste3_rv with an expected band size of 1,136 bp (black arrowhead).

**Supplementary Figure 8:**
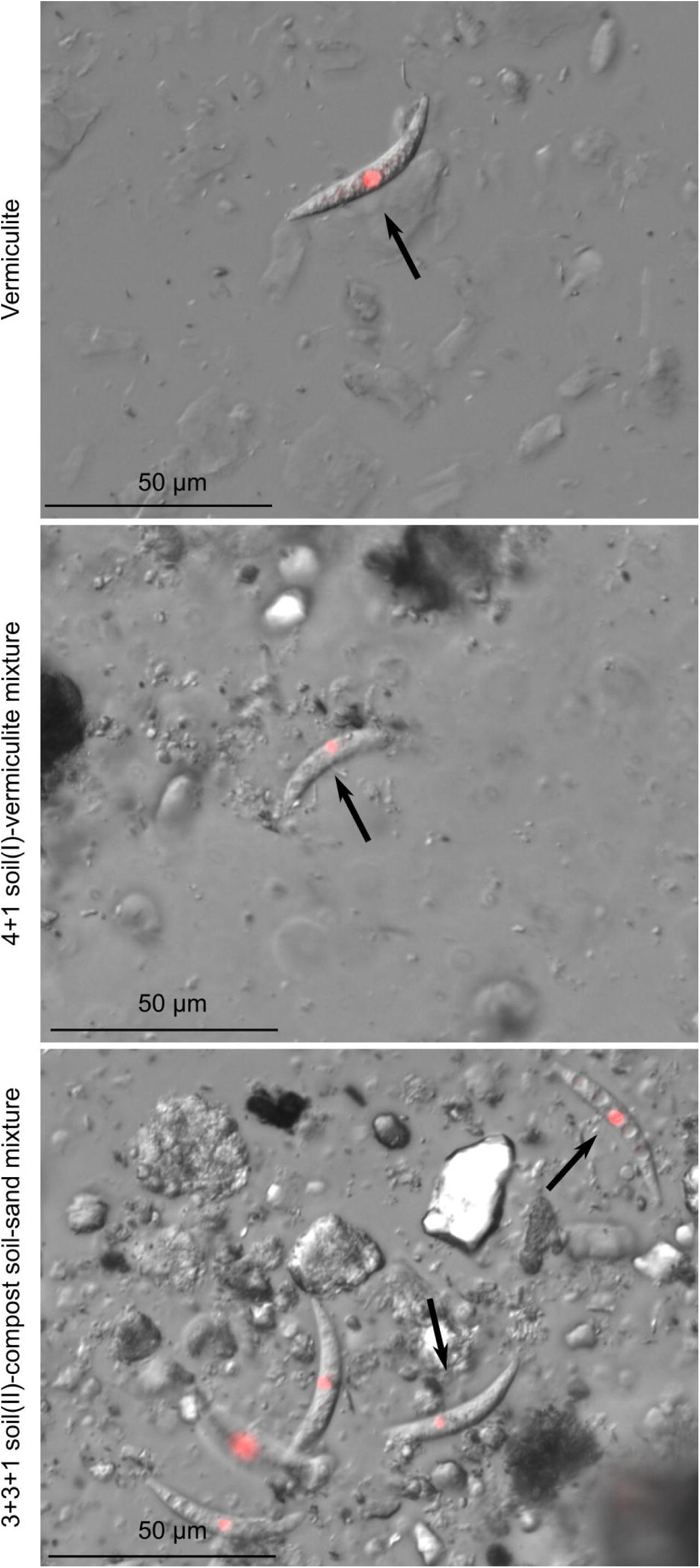
Symptom development after dipping root infection of *Z. mays* with *C. graminicola*. Roots of 5 d old plants were dipped in conidial suspensions. Subsequently, the plants were incubated in vermiculite for 21 d. Quantification of length and biomass of the above ground plant parts. Error bars represent SD calculated from n ≥ 14 experiments, ns = not significant.

**Supplementary Figure 9:**
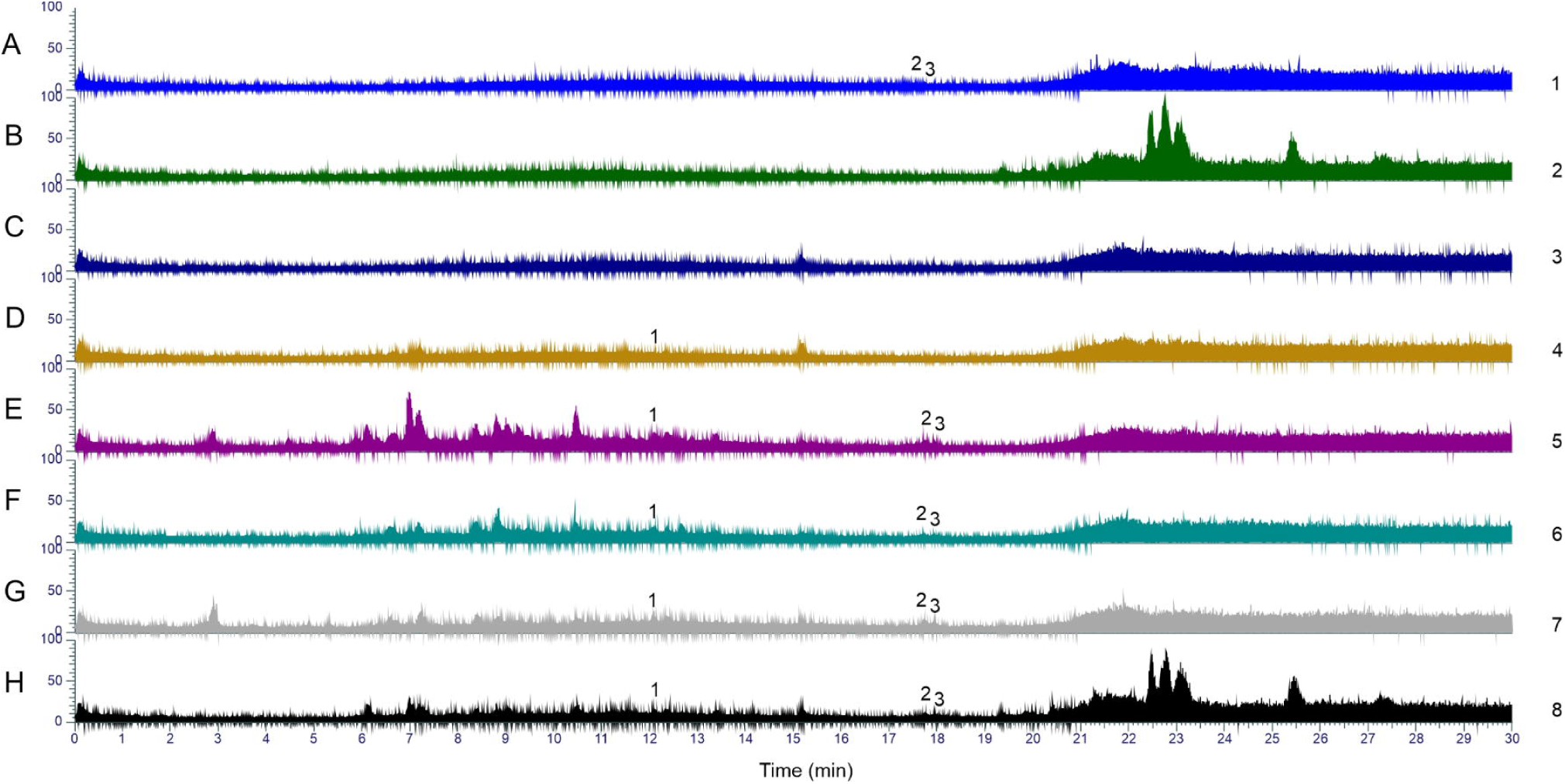
Disease symptoms on a plant co-incubated with oval conidia of CgM2. The plant was grown in conidia enriched vermiculite for 3 weeks. Brown lesions, typical for maize anthracnose, on leaves are indicated with a blue arrow.

**Supplementary Figure 10:**
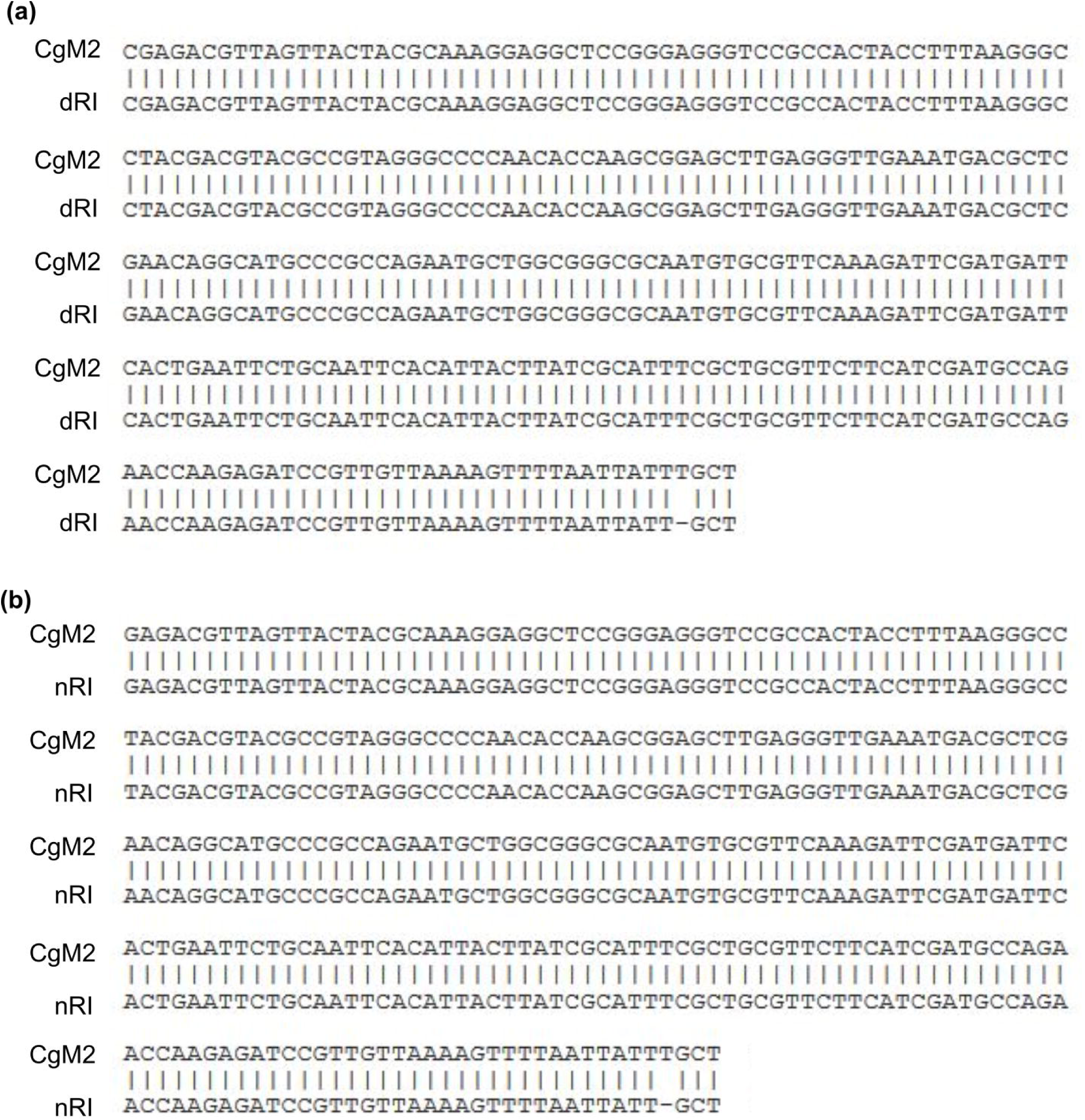
Alignment of CgM2 sequences. CgM2 was aligned with the sequence obtained of a plant stem from the (a) infection by root-dipping (dRI) or (b) infection in conidia enriched vermiculite (nRI). The sequence comprises ITS1, the 5.8S rRNA encoding region and ITS2. Alignments were done with NCBI/BLAST.

Supplementary Table 1 **Oligonucleotides used in this study.**

Supplementary Table 2 ***Colletotrichum graminicola* strains used in this study.**

Supplementary Table 3: **Plasmids used in this study.**

