## Supplementary material for "Maize diterpenoid sensing via Ste3 a-pheromone receptor and rapid germination of *Colletotrichum graminicola* oval conidia facilitating root infection": S Table 2

**Table S2** ***Colletotrichum graminicola strains used in this study.***

| Strain | Genotype | Reference |
|---|---|---|
| CgM2 (M1.001) | *C. graminicola* wild-type (wt) | (Forgey *et al.*, 1978) |
| CgM2::arp1-tagRFP-T | ectopic integration of pCgarp1-TagRFP-T in CgM2; *gen*$^{R}$, ssi, *Cgarp1P::Cgarp1::TagRFP-T::TtrpC* | (Groth *et al.*, 2021) |
| CgM2::RH2B | ectopic integration of Pgpd::hh2b::tdTomato::TtrpC in CgM2; *hyg*$^{R}$, ssi, *CgM2::rh2b* | (Nordzieke *et al.*, 2019) |
| ΔCgste3 | Homologous replacement of *Cgste3* in CgM2, ssi, *hyg*$^{R}$, *Cgste3*::*hph*, | this study |
| ΔCgste3::Cgste3 | Ectopic integration of pCgste3_nat in ΔCgste3, *nat*$^{R}$, ssi; Δ*Cgste3*::*Cgste3* | this study |

*gen*$^{R}$: resistant to geniticin; *nat*$^{R}$: resistant to nourseothricin; *hyg*$^{R}$: hygromycin resistant; ssi: single spore isolate
