## Supplementary material for "Maize diterpenoid sensing via Ste3 a-pheromone receptor and rapid germination of *Colletotrichum graminicola* oval conidia facilitating root infection": S Table 1

**Table S1 Oligonucleotides used in this study.**

| Oligonucleotide | Sequence (5` to 3`) |
| --- | --- |
| ITS_P4_fw | GCCGGAGGATAACCAAACTCTG |
| ITS_P9_rv | GATCCCGATGCGAGACGTTAG |
| GFP-f | ATGGTGAGCAAGGGCGAGGAGC |
| GFP-r | CTTGTACAGCTCGTCCATGCCGAGAGTG |
| hph-f | GTTAACTGATATTGAAGGAGCATTTTTGG |
| hph-r | GTTAACTGGTTCCCGGTCGGCATCTACTC |
| ste3_P_fw | GTAACGCCAGGGTTTTCCCAGTCACGACG<br>CAATTGTACCCCTCTTCCCGTAC |
| ste3_P_rv | GAGTAGATGCCGACCGGGAACCAGTTAAC<br>CTAAGTAAGTGTTCGAAACGACGC |
| ste3_T_fw | CCAAAAATGCTCCTTCAATATCAGTTAAC<br>CTCGCTTTATCCCAGAACGTTG |
| ste3_T_rv | GCGGATAACAATTTCACACAGGAAACAGC<br>GGACGAACCGACATGAATTTATACG |
| ste3_c_fw | CTTCCGGATGGCGAT CAATTGTACCCCTCTTCCCGTAC |
| ste3_c_rv | CCCTGCCCCTGAGAT GGACGAACCGACATGAATTTATACG |
| Ste3_fw | CACAGTACACCAACGCCTATCTC |
| Ste3_rv | CTACTCTGTCTCTACCTGCAGC |

Overhangs for the assembly reactions are indicated in red.
