## Supplementary material for "Maize diterpenoid sensing via Ste3 a-pheromone receptor and rapid germination of *Colletotrichum graminicola* oval conidia facilitating root infection": S Table 3

**Table S3 Plasmids used in this study.**

| Plasmid | Features | Reference |
|---|---|---|
| peGFP-Cgatg8_gen | 5‘ Cgatg8::eGFP::Cgatg8::3’Cgatg8, genR, ampR | (Nordzieke, 2022) |
| pCgste3_KO | *5‘ Cgste3::hph::3’ Cgste3, hyg$^R$, amp*$^R$ | this study |
| pCgste3_nat | *5‘ Cgste3::Cgste3::3‘ Cgste3, nat$^R$, amp$^R$* | this study |
| pJet1.2 | *amp$^R$* | ThermoFisher Scientific |
| pJet_nat | *nat$^R$, amp$^R$* | (Nordzieke, 2022) |

*amp$^R$*: ampicillin resistant; *nat$^R$*: resistant to nourseothricin; *URA3*: encodes for Orotidine-5'-phosphate (OMP) decarboxylase
